# ENvironmental Dynamics Underlying Responsive Extreme Survivors (ENDURES) of Glioblastoma: a Multi-disciplinary Team-based, Multifactorial Analytical Approach

**DOI:** 10.1101/461236

**Authors:** Sandra K. Johnston, Paula Whitmire, Susan Christine Massey, Priya Kumthekar, Alyx B. Porter, Natarajan Raghunand, Luis F. Gonzalez-Cuyar, Maciej M. Mrugala, Andrea Hawkins-Daarud, Pamela R. Jackson, Leland S. Hu, Jann N. Sarkaria, Lei Wang, Robert A. Gatenby, Kathleen M. Egan, Peter Canoll, Kristin R. Swanson, on behalf of the ENDURES consortium

## Abstract

Although glioblastoma is a fatal primary brain cancer with a short median survival of 15 months, a small number of patients survive more than 5 years after diagnosis; they are known as extreme survivors (ES). Due to their rarity, very little is known about what differentiates these outliers from other glioblastoma patients. For the purpose of identifying unknown drivers of extreme survivorship in glioblastoma, we developed the ENDURES consortium (ENvironmental Dynamics Underlying Responsive Extreme Survivors of glioblastoma). This consortium is a multicenter collaborative network of investigators focused on the integration of multiple types of clinical data and the creation of patient-specific models of tumor growth informed by radiographic and histological parameters. Leveraging our combined resources, the goals of the ENDURES consortium are two-fold: (1) to build a curated, searchable, multilayered repository housing clinical and outcome data on a large cohort of ES patients with glioblastoma and (2) to leverage the ENDURES repository for new insights on tumor behavior and novel targets for prolonging survival for all glioblastoma patients. In this article, we review the available literature and discuss what is already known about ES. We then describe the creation of our consortium and some of our preliminary results.

**Funding:** This review was financially supported by a grant from the James S. McDonnell Foundation

**Conflicts of Interest:** The authors have declared that no conflicts of interest exist.

**Authorship:** Conceptualized consortium: LW, RG, KME, PC, and KRS. Built consortium: SKJ, PK, NR, JS, KME, PC, and KRS. Wrote the manuscript: SKJ, PW, SCM, PK, AP, and KME. Reviewed and edited the manuscript: LFGC, MMM, AHD, PRJ, and LSH. Contributed to writing, provided feedback, and approved of final manuscript: All authors.

**Link to website for ENDURES:** http://mathematicalneurooncology.org/?page_id=2125

## Extreme survival among glioblastoma patients

Glioblastoma (GBM) is a notoriously aggressive primary brain cancer and is uniformly fatal, with a median survival of 15 months^1,2^. The use of a multimodal therapeutic approach, involving surgery, radiotherapy, chemotherapy, and tumor-treating fields, aims at prolonging overall survival, but often only increases life span by a few months^3, 4^. Yet, a small percentage of GBM patients survive more than 5 years after diagnosis; these patients represent extreme survivors (ES) of GBM. Identifying factors associated with extreme survivorship could lead to the discovery of new life-extending therapies.

Estimates of 5-year survival rates among newly diagnosed GBM patients vary greatly, depending on the reporting institution, decade of patient treatment, and course of patient treatment. Five-year survival rates estimated before 2005, ranging from 3% to 12%^5–7^, may not be comparable to today’s estimates due to changes in grading criteria and frequent misdiagnosing of initial tissue specimens^8,9^. More recently, 5-year survival rates for patients receiving maximal safe resection, concurrent radiotherapy and chemotherapy, and adjuvant chemotherapy are approximately 10%^3^. This rate is known to vary based on molecular and genetic features and patient age. For example, patients under 50 years of age have a 5-year survival rate of 17%, while patients over 50 have a 5-year survival rate of 6.4%^3^. Establishing accurate 5-year survival rates allows us to compare institutions and treatment regimens and to identify disparities in GBM outcome^10^. In addition to basic survival rate determination, there is a need for conditional prognostic estimates that can be applied throughout the course of a patient’s treatment. For example, Johnson et al. found that the probability of surviving to 5 years was 31% among patients who had already survived 2 years and 57% among patients who had already survived 3 years^11^. While the likelihood of becoming an ES seems low at the time of diagnosis, this outcome becomes increasingly more likely as the patient crosses certain milestones of survival^12^. A detailed understanding of these milestones can generate prognostic information that maintains relevance throughout the disease course.

While it is certainly every patient and clinician’s goal for the GBM patient to become an ES, it is vital to consider the quality of life and neurocognitive function that patients experience at this stage. Studies on quality of life have found that many ES are able to work full-time, live independently, retire, or raise children with minimal neurological deficit^13–15^. However, other ES have to take disability pensions, become dependent on caretakers, or suffer severe neurological deficits^13–15^. It is important to discuss the balance between aggressive long-term treatment plans and patient quality of life, and tracking this information in ES provides the prerequisite data for having this discussion.

Collecting data on large numbers of ES permits researchers to identify potential predictors of ES and possible targets for therapeutic agents. These predictors could be present at any stage of the patient treatment process and could be based on factors like patient demographics or molecular and genetic features. With small sample sizes of less than 15 ES, two studies found that a majority of ES patients presented with seizures^6,16^. Previous literature has also established that long-term survivors (3+ years) tend to be younger than the average GBM patient^16^. Multiple studies found that ES cohorts have an average age of 40 years at time of diagnosis^7,8,13,17^, while the average age of primary GBM patients tends to be above 60 years^18,19^. These studies also observed that ES tend to have higher Karnofsky Performance Status (KPS) scores at time of tumor presentation^6,7,13,20^. Given that GBM is more common in males than females^20,21^, some studies had disproportionately large representation of females in their ES cohorts^6,7,13,20^; larger sample sizes will be needed to statistically validate whether ES is more common among females.

There is minimal research on potentially distinguishing genetic and molecular features of GBM ES and available studies are limited by small numbers of patients. A study of 17 ES found a greater prevalence of intermediate fibrillary elements and a lower prevalence of small anaplastic elements in ES when compared to other GBM patients^8^. Histologically, a study of six ES found that three had features of giant cell GBM, which is a much higher prevalence than the commonly observed rate of 5% among GBM patients overall^20^. Among ten 7+ year survivors, four had overexpressed p53 and four had EGFR overexpression^22^, while a study of 33 ES found that ES had TP53 mutations less frequently than long-term survivors (3+ years)^23^. While there are many studies on the genetic and molecular features common to long-term survivors, our review did not find any additional information on the features common among ES specifically.

Multiple clinical trials and studies have shown the impact of treatment upon the potential for prolonged survival. Stupp et al. observed an 9.8% 5-year survival rate among patients who received radiotherapy and concomitant and adjuvant temozolomide and a 1.9% rate among those who received radiotherapy alone^3^. Other studies have retrospectively observed that ES often received multimodal therapy including chemotherapy and radiotherapy^7,8,17,23^. However, extent of initial surgical resection does not appear to impact 5-year survival rates^3^. After receiving initial therapy, ES often have longer periods of disease stability before recurrence or progression^6,7,13^. One notable study on disease recurrence in ES reported that multiple patients experienced recurrence after 10+ years of stable, progression-free survival (PFS). The author concluded that long PFS is not evidence of cured disease and that vigilant monitoring for radiographic or symptomatic changes is necessary at all points of survival^14^.

The key problem with understanding the features and drivers of extreme survival is that most data sets do not have sufficient numbers of ES patients for study^24^. Therefore, investigation of these rare GBM patients requires multi-center collaborative efforts and a structured sharing environment. The ENDURES (ENvironmental Dynamics Underlying Responsive Extreme Survivors of glioblastoma) consortium (funded by the James S. McDonnell Foundation) collects the data of ES patients from across the country, creating a data set primed for multi-pronged analysis **(Figure 1)**. It is our goal to use this analysis to contribute to the scientific community’s understanding of the drivers of ES and the discovery of actionable targets for drug development and intervention.

**Figure 1.**
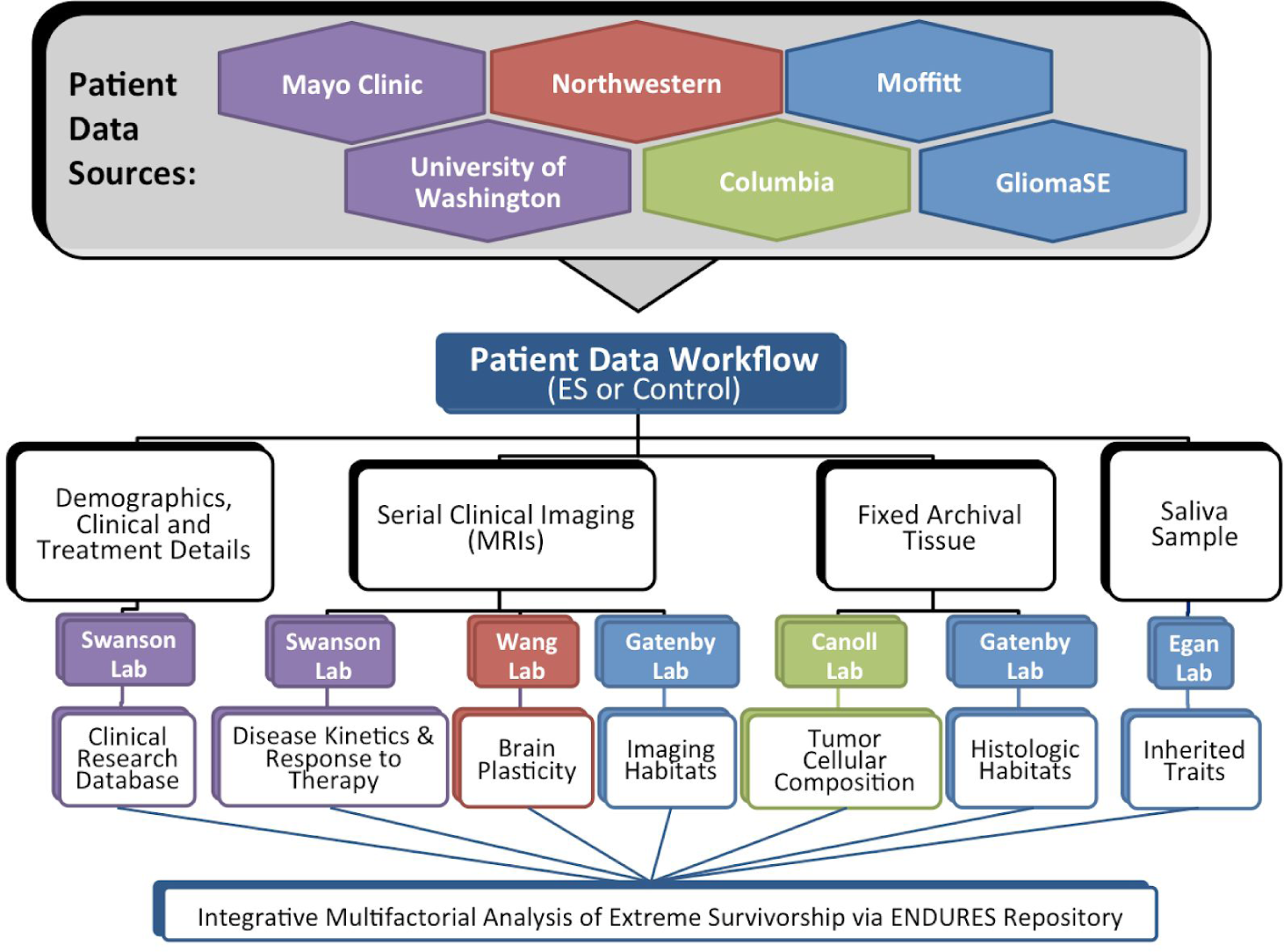
Visual representation of the data collection and analysis workflow within the ENDURES consortium.

## Building a multi-institutional consortium and collecting data on extreme survivors

### Consortium Building

*Core challenge: Creating a large cohort of ES requires combining the patient databases of multiple institutions.* Our consortium currently consists of five institutions: Northwestern University, University of Washington, Moffitt Cancer Center, Columbia University, and the Mayo Clinic. These institutions contributed patients from their internal records (either from a research database or by manually retrieving individual patients from the medical records), public data sets (including TCGA patients), and other collaborative glioma studies (including the GliomaSE study). As of this writing, we have recruited 91 ES patients, both alive and deceased, to the ENDURES database. We also have 36 patients (9 alive and 27 deceased) with prolonged survival of 3-5 years. Growing the consortium remains a priority and partnering with additional institutions will help us achieve this goal. Inquiries regarding expanding or creating new partnership should be directed to corresponding author KRS, and interested institutions should visit http://mathematicalneurooncology.org/?page_id=2125.

### Regulatory Considerations

*Core challenge: Ensuring that each institution’s regulatory agreements allow for data-sharing and acquisition with multiple institutions.* The task of building an anonymized, multi-institutional data-sharing repository can be a regulatory and logistical challenge. Each home institution was responsible for ensuring that its regulatory agreements, including institutional review boards (IRB), permitted the collection of patient data and the sharing of this data with the central collecting institution (Mayo) and entire consortium. At the central collecting institution, the regulatory agreements had to permit the collection and sharing of data from the source institutions, the storage of data in the repository, and the conduct of research and analysis. When available, archived patient tissue was sent to a central pathology institution (Columbia) for storage and evaluation. Data use and transfer agreements were required by the institutions that shared tissue with the central pathology core.

### Patient Identification

*Core challenge: Identifying patients at the contributing institutions and maintaining longitudinal follow-up.* In order for patients to meet the criteria for the ENDURES database: A) GBM had to be the first glioma diagnosis received, B) the GBM diagnosis had to be pathology-confirmed, and C) patients had to have survived at least 5 years after the GBM diagnosis (≥ 1825 days). Each center identified pathology-confirmed, first-diagnosis GBM patients with prolonged survival from their local pre-existing internal records and databases. Patients surviving at least five years after their diagnosis were added to the ENDURES database. Other patients with prolonged survival (3-5 year survival) were considered “potential extreme survivors” and were followed by their neuro-oncologists and our research team to include them in the ENDURES cohort if survival passed the 5 year mark. Once patients were identified, their anonymized data was collected and consolidated for entry into the ENDURES database.

### Clinical Data Acquisition

*Core challenge: Acquiring and entering diversely-formatted clinical data for standardized presentation.* In order to capture sufficient data for an extensive multifactorial investigation, we aimed to amass a comprehensive set of information on each patient **(Figure 2)**. Targeted clinical data included patient age, sex, overall survival (OS), presenting symptoms, performance scores (KPS), and more. For histological information, we collected digitized and/or physical stained tissue slides and available molecular phenotyping. Multi-modal pre-surgical and serial post-surgical (if available) MR images were collected. Relevant treatment dosing and timing information was compiled, which could include surgeries (extent of resection), chemotherapy, radiation therapy, and more. Each center was responsible for entering its own data into the central ENDURES repository. This process was facilitated by the senior clinical data coordinator at the central institution (Mayo), who provided training on data entry and verified incoming data for completeness, accuracy, and consistency. Due to variations in the formatting and completeness of patient records at the home institutions, data entry into the database was a challenging and tedious task. All patient data were coded for anonymization in order to protect patient privacy and follow regulatory requirements.

**Figure 2.**
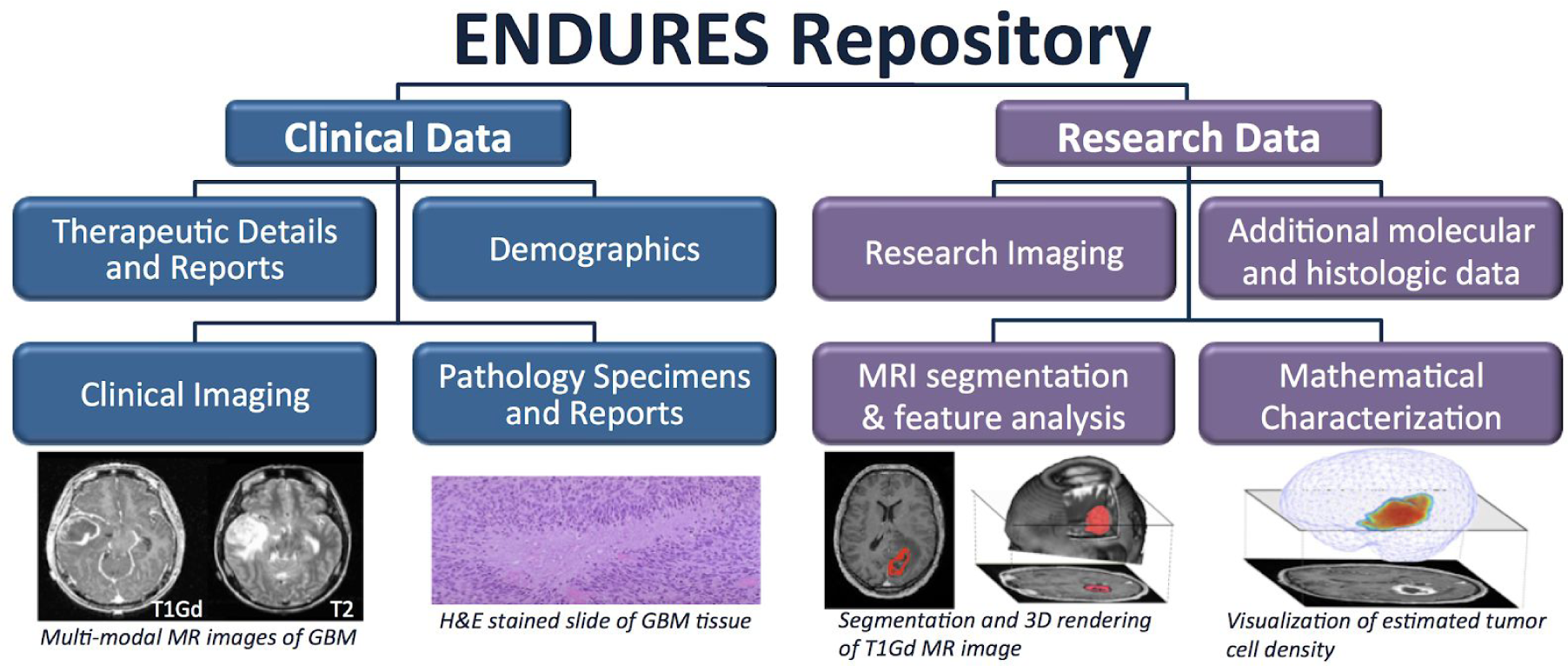
Collecting and processing diverse types of patient data allows for multi-factorial analysis. Blue boxes represent data extracted from the medical records, tumor tissue, and images; purple are generated through our research efforts.

### Data Repository with Longitudinal Follow-up

*Core challenge: Ensuring that the central database is accessible, well-maintained, and current.* An important goal of the ENDURES consortium is to facilitate the sharing of data across sites. For this reason, the central collecting institution (Mayo) had to store the data in a repository that could be accessed remotely by select individuals. We leveraged the existing Swanson Lab Brain Tumor Repository (BTR), which was created in-house with IRB approvals, extending its capabilities as a robust platform for data acquisition, storage, and sharing. The data is managed and accessed via a web-based application of the BTR built using standard web architecture running on infrastructure in a public cloud. The application is built with modern Open Source tools and frameworks to facilitate ease of operation and modification in the future. The BTR can be accessed online, although access is highly controlled. Before entry into the BTR, patient images and information are coded for de-identification and privacy purposes. Despite coming from widely varied home institutional formatting, data in the BTR is represented in a uniform format **(Figure 3).** This important detail is essential for research and allows for querying of the entire dataset.

**Figure 3.**
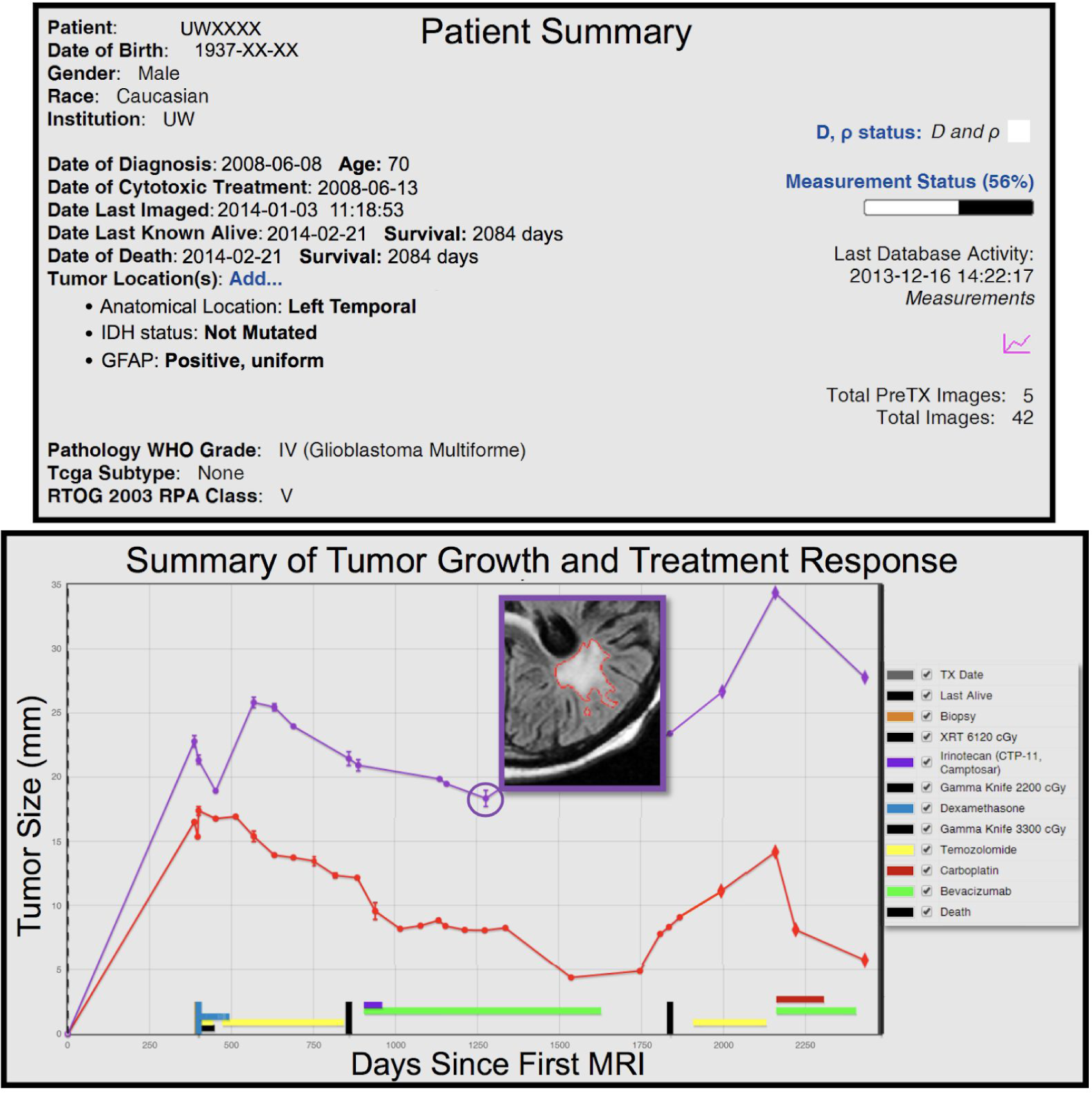
Example of annotated repository data for single patient. Top: Patient summary of key characteristics with dates altered for de-identification. Bottom: Volumetric data gathered over time from segmented, multi-modal MR images and clinical therapies annotated. Pictured is one slice of a segmented T2-weighted fluid-attenuated inversion recovery (FLAIR) MR image.

### Pathology Core

*Core challenge: Acquiring tissue from the initial resection when surgery was completed at an outside institution.* When available, patient tissue (in tissue microarray when possible, otherwise as hematoxylin and eosin stained slides or digitized images) was sent from the home institution to the central pathology institution (Columbia) for analysis. Frequently, institutions needed to keep specimen samples for future diagnostic confirmation, leaving very little tumor tissue available for research. Many of our patients were diagnosed before molecular testing for features such as MGMT methylation and IDH1 mutation were commonplace. Ideally, we would use this stored patient tissue to perform these tests and additional tests for factors like cell lineage markers, cell density, and proliferation markers.

### Multimodal Data Analysis

Collecting such a wide variety of data enables us to perform a variety of analyses to identify determinants of extreme survivorship, including: mathematically modeling tumor kinetics^25–31^, measuring metrics of brain plasticity^32–34^, profiling RNA-expression, contributing to genetic association studies^35–37^, and identifying tumor habitats^38^ **(Figure 1)**. Using patient-specific mathematical models of tumor growth kinetics, we hope to gain insight into GBM growth patterns and assess response to therapy. With MRI analysis, we aim to find image-defined ‘habitats’ associated with extreme survivorship and characterize tumor ecology and evolution using texture analysis and radiomics. In addition to the ES added by the ENDURES consortium, the BTR also contains the MR images and patient data of over 1700 non-extreme surviving GBM patients. These patients can be used as controls and will be compared to ES in the multimodal analysis.

## Preliminary results from analysis of our extreme survivors

Our research group has already completed a number of preliminary analyses on our ES cohort. Using biomathematical models, we found that extreme survivors have lower rates of image-based estimates of tumor cell proliferation than their shorter-surviving counterparts^39^. Additionally, a review of our cohort found the occurrence of giant cell morphology is higher among our ES patients (17.8%) (S.K. Johnston, unpublished data)^40^ than it is among the general GBM population (5%)^20^. A number of investigations have combined ES with other long-term survivors (LTS) (OS > 3 years) and compared them to their shorter-surviving counterparts. One of these investigations found that LTS reached minimum tumor volume later and had a significantly slower rate of post-radiation tumor growth compared to non-LTS (OS < 3 years)^41^. A second investigation comparing LTS to patients with OS < 500 days, found that LTS were younger, had higher initial KPS, and more commonly had cystic tumors^42^. Another study found that among right temporal GBM patients, LTS had greater contralateral cortical thickness than non-LTS^43^. Lastly, tumors in LTS have higher fraction of high contrast enhancement and high FLAIR signal intensity compared to short-term survivors (OS <19 months)^44^.

## Case reports from our data set

In order to demonstrate the diverse disease and treatment courses of ES patients, we have included four case reports below **(Figure 4)**.

**Figure 4.**
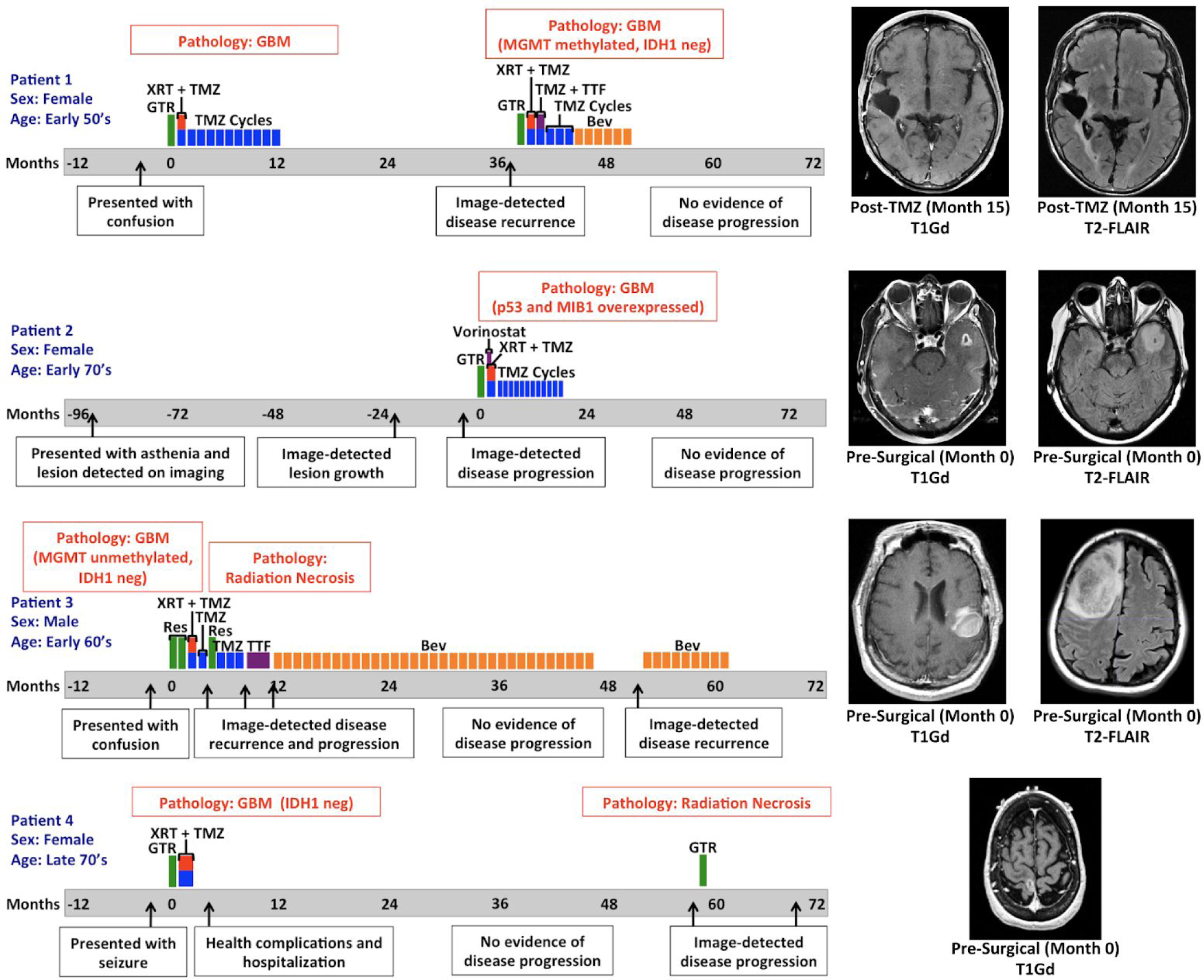
ES Patient Cases: Timeline of disease and treatment course. Time zero represents the date of GBM diagnosis (usually on the date of first resection). Closest available MR images to date of diagnosis are shown. (T1Gd = gadolinium-enhanced T1-weighted, T2-FLAIR = T2 fluid-attenuated inversion recovery, GTR = gross total resection, XRT = radiation therapy, TMZ = temozolomide, MGMT = O(6)-methylguanine-DNA methyltransferase promoter, IDH1 neg = negative for isocitrate dehydrogenase 1 mutation (R132H) by IHC, TTF = tumor treatment fields, Bev = bevacizumab, Res = resection)

Patient 1 is a female in her early 50’s who came to medical attention in the late 2000s because of progressive episodes of confusion. She was found to have a right temporal heterogeneously enhancing mass and ultimately underwent gross total resection. Pathology was consistent with glioblastoma. She completed concurrent radiation therapy and temozolomide without adverse effect, subsequently receiving 10 cycles of adjuvant temozolomide. The patient had evidence of radiographic disease recurrence 38 months after her original date of diagnosis and underwent re-resection where a gross total resection was achieved. Pathology was consistent with recurrent glioblastoma and found to be negative for mutant IDH1 (R132H) by immunohistochemistry (IHC) and MGMT methylated. She was given radiation therapy with concomitant temozolomide and went on to receive 4 adjuvant cycles of temozolomide. Tumor treating fields were implemented during the first cycle of adjuvant temozolomide but then discontinued after one month because of poor tolerance. She then switched to bevacizumab due to concern for post-treatment effect and corticosteroid intolerance. The patient completed 6 cycles of bevacizumab and has remained off treatment since that time. Last neuroimaging continues to have no evidence of active disease, more than 80 months from date of original diagnosis.

Patient 2 is a female in her early 70’s who came to medical attention in early 2000s for occasional episodes of asthenia in her left leg. Ultimately, an MRI of the brain was performed and she was found to have an incidental asymptomatic nonenhancing left temporal lesion. The patient underwent serial imaging which showed interval stability followed by eventual growth. With concern for language deficit from surgical resection, surgery was delayed until the lesion developed a central area of contrast enhancement, which happened 8 years after the initial image finding. A gross total resection removed the enhancing lesion and surrounding FLAIR abnormality. The pathology was consistent with glioblastoma (p53 mutated and elevated MIB 1 index). The patient went on to receive radiation therapy with concomitant and adjuvant temozolomide with vorinostat on clinical trial N0874^45^. The patient tolerated 12 adjuvant cycles of temozolomide without significant adverse effect, but the study drug was discontinued after the first adjuvant cycle because of toxicity. Now over 77 months out from GBM diagnosis, the patient has not received treatment in the last four years and has shown no evidence of disease recurrence.

Patient 3 is a male in his early 60’s who presented in the early 2010s with confusion. He was then found to have an enhancing right frontal mass that was nearly totally resected and diagnosed as glioblastoma (MGMT unmethylated and negative for mutant IDH1 by IHC). After surgery, he underwent a re-resection followed by standard of care radiation therapy with concurrent temozolomide and one cycle of adjuvant temozolomide. One month later, he presented to the clinic with worsening left sided weakness and falls and was found to have an enhancing mass recurring in the right frontal lobe. This mass was resected and found to be necrotic tissue, and adjuvant temozolomide therapy was continued. Two months later, progression was detected and he was started on tumor treatment fields until disease progression was detected again. At this point, the patient agreed to start bevacizumab monotherapy and within 6 weeks of beginning the therapy, the patient showed both symptomatic and radiographic response. He continued on bevacizumab and remained stable for almost three years until month 48. Almost 52 months after the initial diagnosis, progression was once again seen and bevacizumab rechallenge was attempted but aborted due to renal impairment. Patient is alive at the time of this writing, but is no longer receiving treatment.

Patient 4, a female in her late 70’s, presented with a focal left sided motor seizure and was found to have an enhancing mass in the right perirolandic area. She underwent gross total resection of this lesion in the early 2010s and pathology reports indicated it was a glioblastoma negative for mutant IDH1 by IHC (R132H). She received standard of care focal radiation with intensity modulated radiation therapy and concurrent temozolomide for six weeks. After radiation, she suffered a severe infection and other complications that required prolonged hospitalization. Ultimately, the patient made the decision to be monitored off therapy and no adjuvant temozolomide was administered. She continued with surveillance neuroimaging alone until radiographic progression was seen 58 months after initial diagnosis in the same area of her original tumor. She then underwent a gross total resection of this lesion and pathology showed radiation necrosis and treatment effect alone without active tumor. She was again monitored off therapy and over a year later gradual radiographic progression was again seen. This was also associated with clinical decline and worsening left sided weakness. At this time, patient declined further treatment.

## Conclusion

Our collaborative consortium has produced a first-of-its-kind, growing dataset for research on GBM extreme survivorship. We hope that the success and opportunities associated with this consortium will encourage other institutions to create similar data-sharing collaborations. The knowledge gained from this ongoing study will contribute to a better overall understanding of GBM. The comprehensive characterization of the molecular and physiological determinants of extreme survival could lead to the development of an optimized treatment strategy for each GBM patient.

## Acknowledgements

We would like to thank all of the students, clinicians, researchers, and staff members who contributed to the building of the ENDURES consortium and patient dataset, including: Dr. Joshua Jacobs, Dr. Cristina Valencia Sanchez, Dr. Danielle M. Burgenske, Kamala Clark-Swanson, Scott Whitmire, Cassandra R. Rickertsen, Gustavo De Leon, Spencer Bayless, and the undergraduate students on our Image Analysis Team at Mayo Clinic; Noah Peeri, Carrie M. Rozmeski, and Melissa H. Madden at Moffitt Cancer Center; Dr. Alfred Rademaker, Dr. Qinwen Mao, Dr. C. Paula Lewis-de los Angeles, David Corwin, and Jessica Quaggin-Smith at Northwestern University; Dr. Jeffrey N. Bruce and Christina Corpuz at Columbia University; and Dr. Russell Rockne at City of Hope.

